# Indistinguishability domains of neural microcircuit motifs mapped through classification scores of postsynaptic spike counts

**DOI:** 10.1101/2025.09.19.673918

**Authors:** Anjali Naveen Kumar, Raghunathan Ramakrishnan

## Abstract

In the kinetic modeling of a chemical synapse, the reversal potential (*E*_syn_) and conductance amplitude (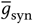) of the synapse jointly regulate the spike-induced activity of the postsynaptic neuron. We simulate two-neuron microcircuit motifs of feedforward (*i* → *j* ) and feedback (*i* ⇌ *j* ) types using random external current pulses provided only at the presynaptic neuron. For combinations of *E*_syn_ and 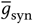, corresponding to an excitatory synapse, a supervised machine learning method for classification distinguishes the motifs perfectly with a score of 1.0 when using binned counts of postsynaptic spikes as the input. Strongly inhibitory combinations of these parameters result in no postsynaptic response in both types of microcircuit motifs; hence, the classifier does not improve upon a random assignment with a score of 0.5. In other domains of the parameter space spanned by *E*_syn_ and 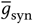, corresponding to diminished excitation/inhibition, the classifier fails (0.5 < score < 1), indicating the challenge in identifying the nature of the synapse inferred exclusively from postsynaptic spikes. For this task, the gradient boosting classifier gives higher classification accuracies that improve with further training, compared to other classifiers explored in this study.

## 1 Introduction

The human brain comprises a highly complex yet coordinated network of over 10 billion neurons [1]. Neurons possess the unique ability to conduct electrical signals through the generation of action potentials, forming the foundation of neural communication. Synapses serve as the junctions between neurons, enabling this communication. Upon stimulation, the presynaptic neuron releases neurotransmitters, which can lead to either excitation or inhibition of the postsynaptic neuron, depending on the type of neurotransmitter molecules (e.g., glutamate, Glu, for excitation and *γ* aminobutyric acid, GABA, for inhibition). These neurotransmitters bind to specific receptors on the postsynaptic neuronal membrane, producing a transient shift in the resting membrane potential (*V*_rest_). This change manifests as either an excitatory postsynaptic potential (EPSP) or an inhibitory postsynaptic potential (IPSP), determined by the type of ion channel activated [2]. EPSPs arise when Glu receptors open [3], allowing Na^+^ and Ca^2+^ ions to enter the cell. This influx drives the synaptic reversal potential (*E*_syn_) toward ∼0 mV, resulting in depolarization [4]. In contrast, activation of postsynaptic ionotropic GABA_*A*_ receptors permits Cl^−^ influx in mature neurons, shifting *E*_syn_ toward approximately −70 mV [5], thereby producing hyperpolarization [6, 7]. While potassium currents play a central role in action-potential repolarization and intrinsic membrane excitability, fast chemical synaptic transmission is primarily mediated by ligand-gated Na^+^, Ca^2+^, and Cl^−^ conductances. In general, if *E*_syn_ lies below *V*_rest_, the postsynaptic effect is inhibitory. On the other hand, if *E*_syn_ exceeds *V*_rest_, the effect is excitatory, provided that the resulting depolarization is sufficient to drive the membrane potential beyond the threshold for spike initiation [8].

The electrical activity of the neuronal membrane determines the impact of synaptic input currents. The driving force of a synaptic current is given by the difference between *V*_rest_ and *E*_syn_ [9]. As current flows, the membrane potential shifts toward *E*_syn_, gradually reducing the driving force. Thus, EPSPs from active synapses do not simply add linearly [9]. Moreover, an increase in synaptic conductance lowers the membrane resistance, which reduces the amplitude and accelerates the decay of EPSPs [9]. Neurotransmitters mediate communication between presynaptic and postsynaptic neurons. Excitatory neurotransmitters, glutamate, act on AMPA/kainate and NMDA receptors, both with reversal potentials near 0 mV. Inhibitory GABAergic synapses act primarily via GABA_*A*_ receptors (*E*_syn_ ≈ −70 mV), with GABA_*B*_ receptors operating at even more hyperpolarized levels (*E*_syn_ ≈ −95 mV) [10, 11]. The exact value of the reversal potential is not fixed but depends on developmental, spatial, and microenvironmental factors. For instance, in immature neurons, intracellular Cl^−^ concentration is higher due to the activity of the Na–K–2Cl cotransporter (NKCC) [12]. As a result, activation of GABA_*A*_ receptors produces depolarization rather than hyperpolarization [7]. With a resting potential of around −75 mV, the opening of relatively few GABA_*A*_ channels can generate large depolarizations in immature neurons due to their high input resistance [12]. Additionally, GABA can reduce excitatory glutamatergic EPSPs through shunting inhibition, the extent of which depends on the density of glutamatergic inputs [12]. During development, this balance shifts: GABAergic transmission transitions from excitatory to inhibitory as chloride homeostasis matures and shunting inhibition becomes dominant [11].

At later stages of development, upregulation of the K–Cl co-transporter isoform 2 (KCC2) drives Cl^−^ efflux, which reverses the transmembrane Cl^−^ gradient and shifts the sign of *E*_syn_. Consequently, activation of the GABA receptor results in influx of Cl^−^, thereby establishing the inhibitory action of GABA [13]. In vitro studies indicate that this upregulation is promoted by “miniature postsynaptic currents”, which occur independently of action potentials. These depolarizations trigger Ca^2+^ signaling, which in turn enhances the expression of the K-Cl co-transporter, KCC2 [12, 14]. This developmental “GABA shift”, together with the maturation of glutamatergic neurons, synapse formation, and experience-driven pruning, shapes a synchronized neuronal network [15]. Balanced excitatory and inhibitory inputs at distinct spatial domains ensure plasticity, synchrony, and proper brain oscillations. For instance, exogenous application of GABA has been shown to inhibit action potentials in pyramidal neurons when applied at the soma but to produce depolarization when applied near dendrites [16]. In hippocampal CA1 pyramidal neurons, GABA can also evoke a biphasic response: an initial hyperpolarization followed by slow depolarization [17]. GABAergic interneurons play central roles in establishing inhibitory circuits essential for orchestrating complex network activity. Disruptions in their development can contribute to neurodevelopmental disorders[18]. For example, in schizophrenia, expression of GAD67, the enzyme converting glutamate to GABA, is reduced in parvalbumin-positive (PV^+^) interneurons, impairing gamma oscillations in the cortex and hippocampus, weakening inhibitory function [19, 20]. Similarly, loss of hippocampal GABAergic interneurons reduces inhibitory control over principal cells, facilitating hyperexcitability and seizure spread in epilepsy [21].

Excitatory and inhibitory synapses are classical paradigms of neuronal communication. Yet, in more complex contexts—such as brain development, where GABAergic transmission initially acts excitatory before shifting to inhibitory, or in epilepsy, where disrupted excitation–inhibition balance drives seizures, this dichotomy becomes less distinct. In this paper, we aim to construct a conceptual map of synaptic behavior in the parameter space defined by *E*_syn_ and *g*_syn_. By simulating prototypical cortical fast-spiking microcircuit motifs and quantifying synaptic effects using a classification score from a supervised machine learning (ML) method, we identify regions that correspond to distinct synaptic regimes.

This paper is organized as follows. In Section 2, we introduce the mathematical models used to simulate current-pulse stimulation on neural microcircuit motifs and describe the two motifs analyzed in this work. Section 3 presents the classification results, where binned spike counts of postsynaptic neurons are used as features across different values of *E*_syn_ and 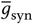, capturing excitatory, inhibitory, and intermediate synaptic regimes with reduced polarity. We examine the performance of machine learning methods, including improvements in classification scores with increasing training set size, and assess the tradeoff between information content and noise when spike counts are binned at different temporal resolutions. We then identify regions of the parameter space that are classifiable in a data-driven manner, as well as indistinguishable regions where classification fails and intermediate scores emerge. Finally, we conclude by outlining the broader scope of this framework for identifying neural microcircuit motifs.

## 2 Kinetic Modeling of Neuronal Membrane Potential

We simulate the current-pulse dynamics of microcircuit motifs consisting of two neurons, *i* and *j*, connected via a synaptic contact, as illustrated in Figure 1. The dynamics of this coupled system is described by an eight-dimensional Hodgkin–Huxley-type model:

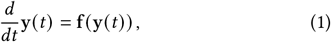

where

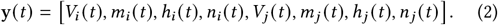

**Figure 1.**
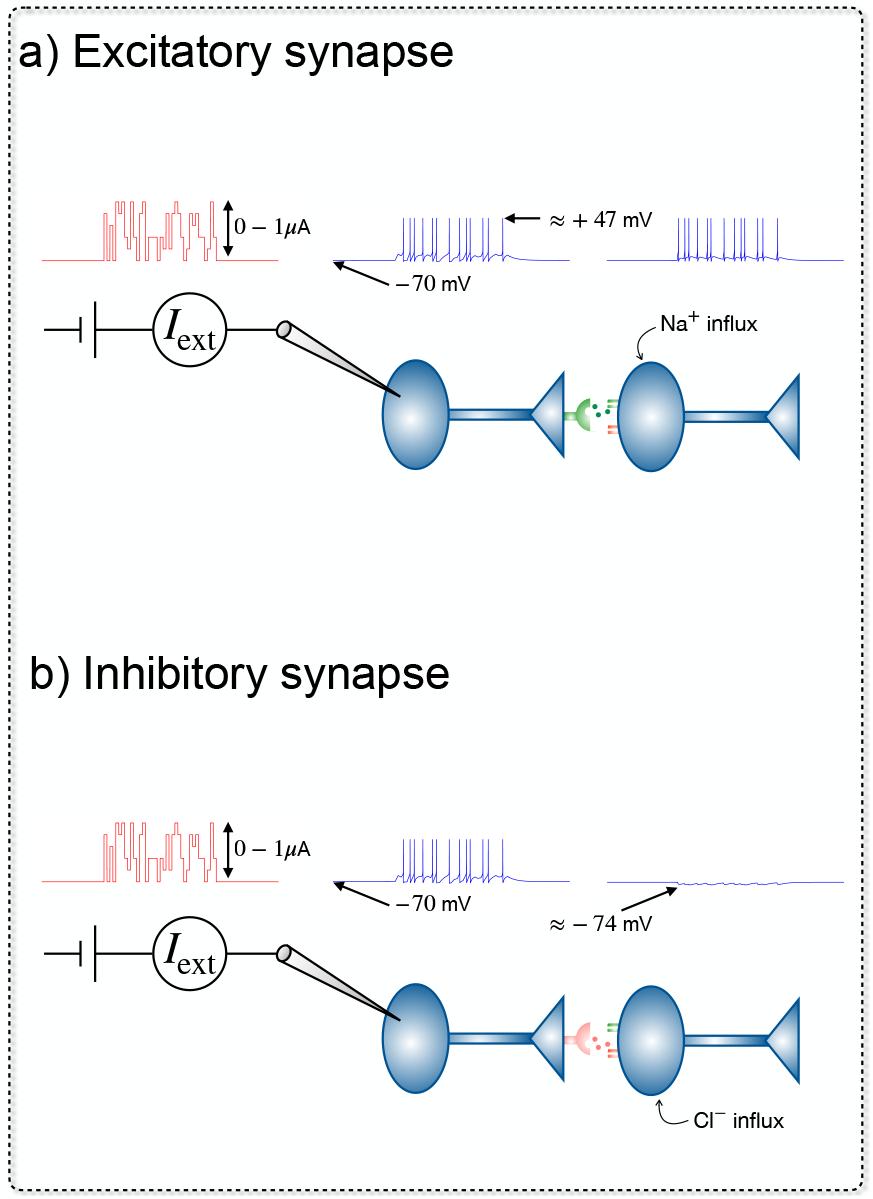
Schematic of excitatory and inhibitory synaptic transmission with identical peak conductance 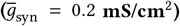: a) Release of neurotransmitters (e.g. glutamate) leading to opening of postsynaptic receptors (ion channels), allowing Na^+^ influx and depolarization toward the excitatory reversal potential (*E*_syn_ = −30 mV); b) Release of neurotransmitters (e.g. GABA) activating ion channels that allow Cl^−^ influx in the postsynaptic neuron and hyperpolarization toward the inhibitory reversal potential (*E*_syn_ = −80 mV).

Here, *V* (*t* ) denotes the membrane potential, and *m*(*t*), *h*(*t*), and *n*(*t* ) represent the gating variables. Each component of

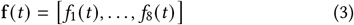

specifies the corresponding differential equation governing one of the eight dynamical variables.

The membrane potential of the presynaptic neuron (*i*) is modeled using the Hodgkin–Huxley framework [22–25]:

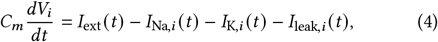

where *I*_Na_ and *I*_K_ represent the sodium and potassium currents that underlie the action potential. Although the classical Hodgkin– Huxley formalism primarily focuses on Na^+^ and K^+^ currents, calcium ions also play a crucial role in neuronal signaling. Mammalian neurons express multiple types of voltage-gated Ca^2+^ channels that permit Ca^2+^ entry, hence, serving as a potent intracellular messenger. Their direct contribution to action-potential upstroke is typically small compared to Na^+^ because Ca^2+^ channels activate more slowly. However, during the peak of the action potential, the depolarization is sufficient to open these channels, allowing Ca^2+^ influx. This influx can in turn activate large-conductance Ca^2+^-activated potassium channels (BK channels or big potassium channels), which drive substantial K^+^ efflux, overpowering the Ca^2+^ entry and accelerating membrane repolarization [26].

Sodium and potassium currents are expressed in terms of the maximal conductances 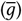, activation variables (*m* for Na^+^ and *n* for K^+^), and the Na^+^ inactivation variable (*h*):

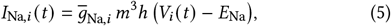

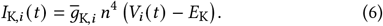

In standard neurophysiological electrode recordings, the neuron effectively acts as a single branch of an electrical circuit. The conductance parameters *g* represent the ease with which ions flow across the membrane, functioning analogously to variable conductances in an electrical network. The direction of ionic flow is determined by the reversal potentials *E*_Na_ and *E*_K_; when *V*_*i*_ (*t*) equals these values, there is no net flux of the respective ions. The leak current represents passive ionic fluxes not explicitly captured by the sodium or potassium channels, effectively accounting for background contributions such as chloride flow [24]:

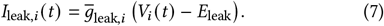

We model two types of microcircuit motifs, as illustrated in Figure 2. For both cases, the spike-driven dynamics of the postsynaptic membrane potential are described by a modified Hodgkin–Huxley model (HHM) without external current:

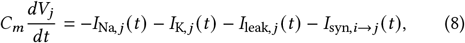

where the notation *a* → *b* indicates that *b* is the postsynaptic neuron receiving input from the presynaptic neuron *a*. The synaptic current is defined as [27]

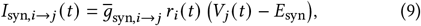

where *r*_*i*_ (*t* ) is the fraction of bound postsynaptic receptors. Initially, *r*_*i*_ (0) = 0, and its dynamics follow [22, 27]

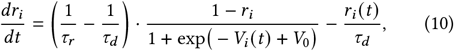

with *τ*_*r*_ and *τ*_*d*_ denoting the rise and decay time constants, respectively. The values of *τ*_*r*_, *τ*_*d*_, and *V*_0_, taken from [22], are listed in Table 1.

**Figure 2.**
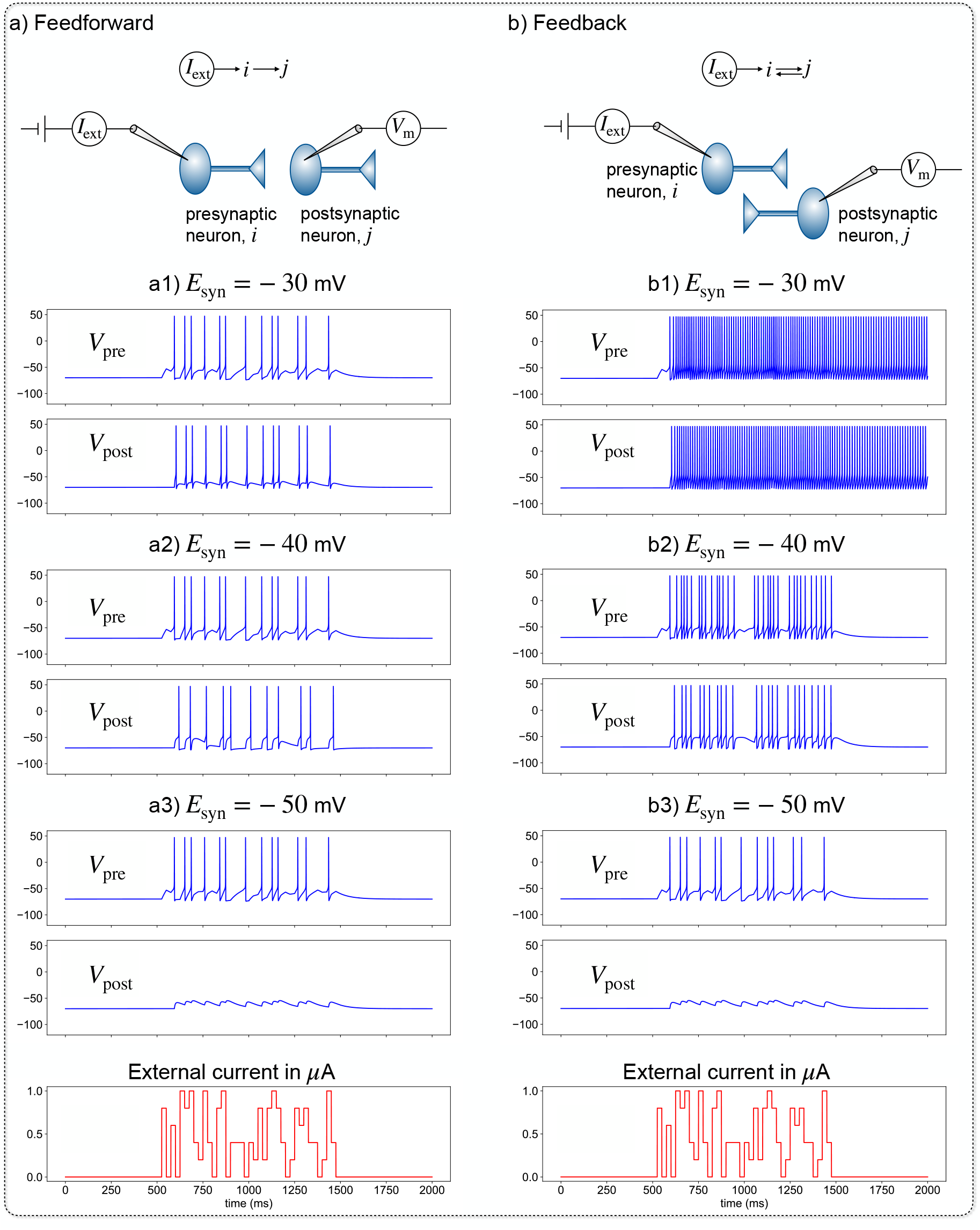
Model two-neuron microcircuits considered in this study: a) Feedforward circuit with neuronal dynamics described by Eq. 4 and Eq. 8; b) Feedback circuit described by Eq. 11 and Eq. 8. For the current pulse shown at the bottom, excitatory dynamics are described by synaptic activity shown in panels a1 and b1, while inhibitory dynamics are shown in panels a3 and b3.

**Table 1.** Parameters employed in solving Eqs.(4, 8, and 11), which describe models of fast-spiking cortical neurons adapted from Ref.[22].

| Model parameter | Unit | Values |
| --- | --- | --- |
| $C_m$ | $\mu\text{F}/\text{cm}^2$ | 1 |
| $E_K$ | mV | -90.0 |
| $E_{\text{Na}}$ | mV | 50.0 |
| $E_{\text{leak}}$ | mV | -70.0 |
| $g_K$ | $\text{mS}/\text{cm}^2$ | 10.0 |
| $g_{\text{Na}}$ | $\text{mS}/\text{cm}^2$ | 56.0 |
| $g_{\text{leak}}$ | $\text{mS}/\text{cm}^2$ | 0.015 |
| $V_T$ | mV | -56.2 |
| $\tau_r$ | msec | 0.5 |
| $\tau_d$ | msec | 8 |
| $V_0$ | mV | -20 |

The difference in the dynamics of both motifs arises from the presynaptic response. For the type-a motif resulting in feedforward communication (see Figure 2 a), the presynaptic potential follows Eq. 4. For the type-b motif (see Figure 2 b), the potential for the presynaptic neuron also includes the synaptic current term as a feedback

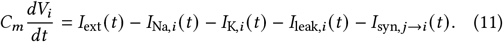

For an excitatory synapse, *I*_syn,*j i*_ *t* can result in perpetual (or recurrent) spikes as shown in Figure 2 (see b1).

The rate equations for the gating variables, common to both preand postsynaptic neurons and adopted from Ref. [22], are given as follows. For the sodium activation variable *m*(*t*)

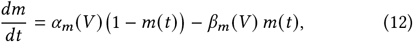

with

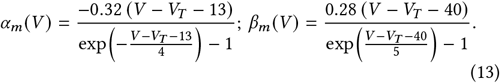

For the sodium inactivation variable *h*(*t*), we have

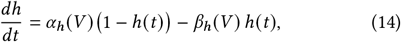

with

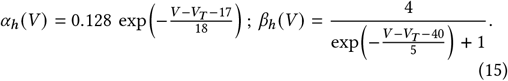

Lastly, the potassium activation variable *n*(*t* ) follows

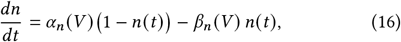

with

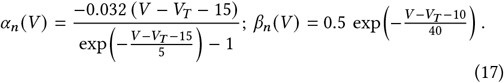

The parameters listed in Table 1 are adopted from Ref. [22]. In that study, fast-spiking (FS) cortical neurons were modeled within the Hodgkin–Huxley framework using only *I*_Na_ and *I*_K_ currents, consistent with our approach. FS neurons are a subtype of cortical interneurons characterized by rapid repolarization, brief action potentials, and the ability to generate high-frequency trains of spikes upon activation [23, 28]. Accordingly, the equations and parameter values employed here correspond to the FS cells of the rat somatosensory cortex.

In our simulations, *V*_rest_ (i.e., *V* (0) of preand post-synaptic neurons) at the initial time (*t* = 0) was set to −70 mV. The driving force of the synaptic current is given by the difference between *V* (*t*) and *E*_syn_. If *E*_syn_ exceeds the membrane potential, the response is excitatory (Figure 1a); and, if *E*_syn_ lies below the membrane potential, the postsynaptic response is inhibitory (Figure 1b). The maximal synaptic conductance 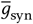 was set to 0.2 mS/cm^2^, consistent with the previous study [22]. The total simulation time was 2000 ms (2 s), with an external current *I*_ext_ (*t*) applied between 500 and 1500 ms. During this interval, for every 25 ms window, a constant current value was randomly selected from the set {0.0, 0.2, 0.4, 0.6, 0.8, 1.0} *μ*A. The effect of varying current-pulse magnitudes is analyzed in Figure 3.

**Figure 3.**
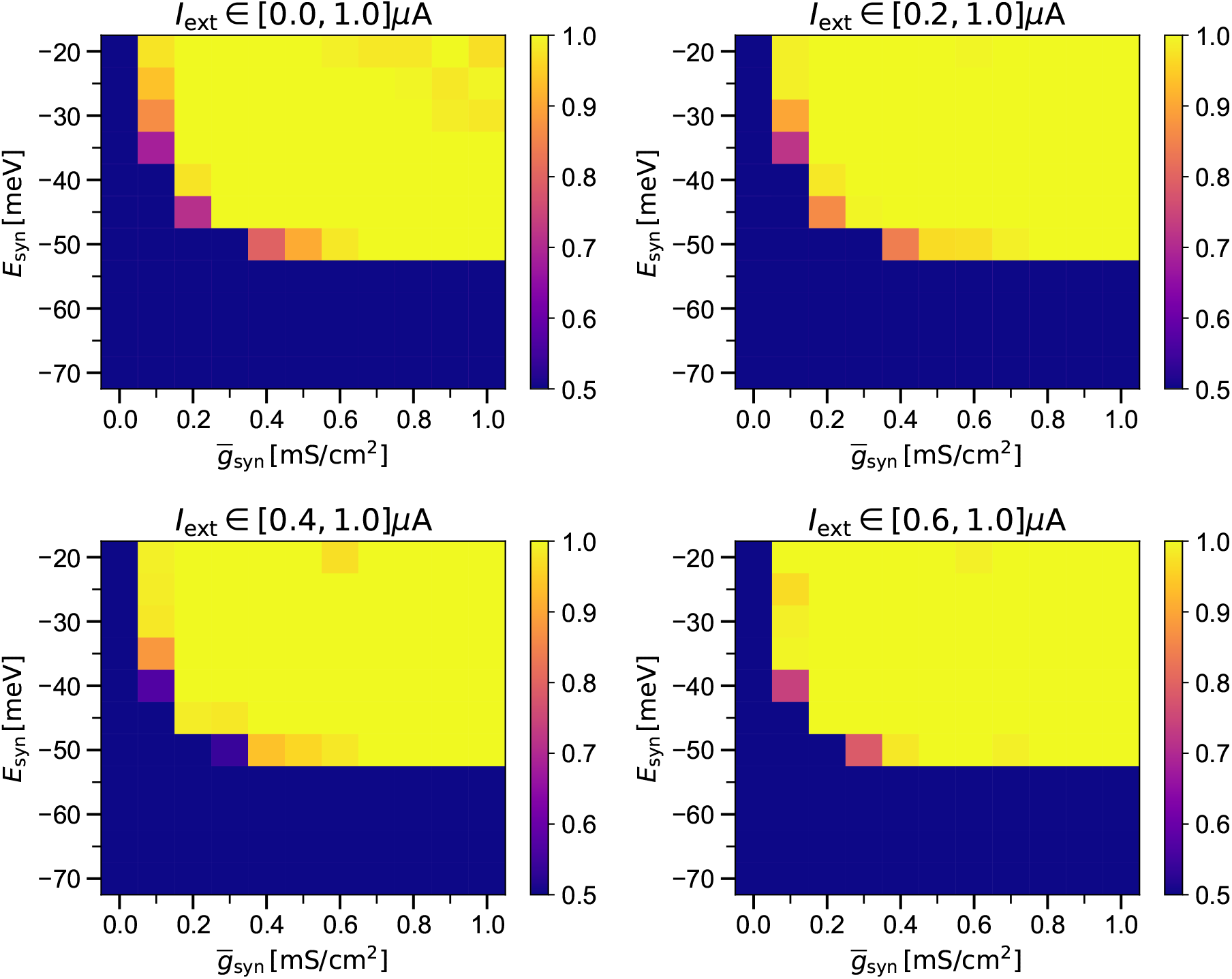
Accuracy, as defined in Eq. 18, for a GB-classifier to identify the microcircuit motifs (see Figure 2) using binned postsynaptic spike counts (total duration: 1000 ms, bin width: 50 ms). Average classification scores are reported as heatmaps for each combination of 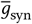 and *E*_syn_ parameters using 5-fold cross-validation of data obtained from simulations of each microcircuit motif using 100 random current pulses of magnitudes specified above the heatmaps.

In clinical neurophysiology, electrical signals must be sampled at sufficiently high temporal resolution to resolve action-potential waveforms. Typical spike durations range from 10^−1^ to 10^−2^ ms, amounting to sampling rates on the order of 10^5^ samples per second. Accordingly, a 1-s recording epoch requires approximately 10^5^ data points, corresponding to a time step of about 0.01 ms [29].

To integrate Eq. 1, we employed the `odeint` solver from the `scipy` library [30], which wraps the well-established *Livermore Solver for Ordinary Differential Equations with Automatic method switching* (LSODA) algorithm [31]. LSODA automatically switches between non-stiff and stiff integration methods and adaptively adjusts the internal step size to preserve numerical stability and accuracy. The tolerances `rtol`=10^−6^ and `atol`=10^−9^ were chosen to control the global error of the solution and to ensure that the integration dynamically accounts for rapid variations in the solution. The maximum allowable step size was limited by setting hmax to 0.05.

Microcircuit motifs form the backbone of neural networks. Two fundamental types are the feedforward and feedback motifs (see Figure 2). In a feedforward motif, neurons are connected in series such that information flows in a single direction [32]. In contrast, in a feedback motif, the presynaptic neuron also receives input from its postsynaptic partner, in addition to external current, which can lead to sustained reciprocal activation. Motifs can be excitatory or inhibitory depending on the neurotransmitter released at the synapse. In modeling, this effect is captured implicitly through the parameters 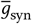 and *E*_syn_.

In addition to feedforward and feedback, other types of motifs are also present in neural networks. One example is lateral excitation or inhibition, where signaling is not transmitted through direct point-to-point synaptic contacts but rather via the diffusion of neurotransmitters, monoamines, or neuromodulators within a population of axons [32, 33]. Another example is disinhibition, a motif in which inhibition is transiently and selectively blocked, thereby producing net excitation [34].

In this study, we analyzed feedforward and feedback motifs under both excitatory and inhibitory regimes by observing the electrical activity of two coupled neurons (Figure 2). In the feedforward case, when an external current is applied to the presynaptic neuron and the synaptic reversal potential is set to *E*_syn_ = −30 mV, the postsynaptic neuron is strongly excited. This occurs because the *V*_rest_ in our simulations is −70 mV, creating a large driving force for the synaptic current. Such values of *E*_syn_ are characteristic of excitatory neurotransmitters such as glu. Under these conditions, the presynaptic and postsynaptic neurons exhibit similar spiking patterns (Figure 2a1). A comparable but more pronounced effect is observed in the feedback (recurrent) motif, where excitation of the presynaptic neuron drives the postsynaptic neuron, which in turn feeds back onto the presynaptic neuron. This recurrent coupling leads to sustained mutual activation and closely matched spike trains in both neurons (Figure 2b1).

When *E*_syn_ is reduced to −40 mV in the feedforward motif, the postsynaptic neuron generates fewer spikes than the presynaptic neuron (Figure 2a2). This reduction reflects the weaker driving force at the lower *E*_syn_. A similar decrease in postsynaptic spiking is observed in the feedback motif (Figure 2b2). However, comparing presynaptic activity across motifs reveals higher spike counts in the feedback case. This is because postsynaptic excitation in the feedback loop reactivates the presynaptic neuron, thereby amplifying its firing.

At *E*_syn_ = −50 mV, the postsynaptic neuron exhibits inhibited spiking activity (Figures 2a3,b3), as synaptic activation remains insufficient to drive the membrane dynamics into the spiking regime. This regime corresponds to shunting inhibition, in which activation of the synapse increases the instantaneous membrane conductance, causing depolarizing current to leak out. At *E*_syn_ = *V*_rest_ = −70 mV, synaptic activation produces no net postsynaptic potential at rest. Further reductions in *E*_syn_ (i.e., *E*_syn_ *<* −70 mV) lead to progressively stronger hyperpolarization as the membrane potential is driven toward *E*_syn_, a hallmark of strongly inhibitory synaptic transmission (Figure 1).

## 3 Classification of Microcircuit Motifs using Supervised Machine Learning

From the current-pulse simulations of both microcircuit motifs, we extracted postsynaptic spike counts and organized them into feature vectors by binning over different time intervals (*N*_bin_). For *N*_bin_ = 1, the feature vector corresponds to the total spike count during the 500–1500 ms window in which the external current was applied. For *N*_bin_ = 2, 5, 10, 20, 40, spikes were summed over bin widths of 500, 200, 100, 50, and 25 ms, respectively. These binned spike counts serve as input features, while the motif labels (0 or 1) are the targets to train supervised machine learning models. The classifiers considered in this study are: K-nearest neighbors (KNN, *k* = 5), stochastic gradient descent (SGD), random forest (RF), support vector classifier with a radial basis function kernel (SVC-RBF), and gradient boosting (GB). All ML calculations were performed using the `scikit-learn` library [35].

For an initial assessment of the performance of various classifiers, we generated a dataset based on simulations with fixed values 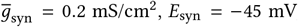, and external current pulses of constant amplitude applied in 25 ms intervals during the 500– 1500 ms window of a total 2000 ms simulation. Current amplitudes were randomly selected from the range [0, 1] *μ*A in steps of 0.2 *μ*A. For each microcircuit motif, 1000 simulations were performed by numerically solving Eqs. (4, 8, 11), with initial membrane potentials (i.e., *V*_rest_) set to *V* (0) = −70 mV. This procedure yielded 2000 data points in total, each consisting of a feature vector (binned spike counts) and an associated target label (0 or 1).

Overall classification accuracy is defined as the fraction of correctly predicted labels over the total number of predictions. In terms of the confusion matrix entries,

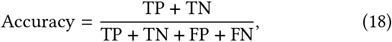

where TP, TN, FP, and FN denote the number of true positives, true negatives, false positives, and false negatives, respectively. All four terms are summed over both classes. The scores reported correspond to the mean accuracies obtained from 5-fold cross-validation. In this procedure, the dataset is partitioned into five subsets of equal size (20% each). Each subset is used once as the test set, while the remaining 80% of the data was used for training. With sufficient training data, the expected minimum classification score is 0.5, which corresponds to random guessing of labels. The upper bound is 1.0, representing perfect classification on the test set. Scores below 0.5 may arise when training sets are minimal, reflecting high variance in model performance due to sampling bias.

Table 2 reports the classification accuracies (as defined in Eq. 18) for all five models considered in this study, using feature vectors constructed from spike counts binned at different temporal resolutions. For training, 80% of the 2000 data points (1000 current-pulse simulations per motif) were used with 5-fold cross-validation. We focused on *E*_syn_ = −45 mV (with 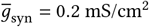), which represents an intermediate regime where the synapse is neither strongly excitatory (*E*_syn_ ≫ *V* (0) ) nor strongly inhibitory (*E*_syn_ ≪ *V* (0) ). In general, ensemble methods such as RF and GB outperform simpler models like KNN and SGD, while SVC (RBF) performs competitively. Cross-validated accuracies in the range of 0.75–0.81 (for *N*_bin_ = 10 in Table 2) across classifiers indicate that spike-binned features provide moderate discriminative power. Performance consistently exceeded the chance level of 0.5, confirming that motif types can be distinguished from spike statistics, though with substantial feature overlap that prevents near-perfect classification. An interesting trend observed across all classifiers is that the maximum accuracy was not obtained at the highest number of bins (*N*_bin_ = 40). This suggests that increasing the temporal partitioning of spike counts does not always add useful information, and at finer bin widths, the classifiers may instead be affected by noise. The highest accuracy of 0.806 was achieved by GB at *N*_bin_ = 10 and *N*_bin_ = 20, showing that although the motifs are distinguishable, the overlap of spike-based features still limits classification performance to well below perfect accuracy.

**Table 2.** Classification scores for various machine learning methods for predicting the labels of two microcircuit motifs using spike counts of postsynaptic action potential from 1000 simulations for each motif. The total simulation time was 2000 ms, and the external current was applied to the presynaptic neuron from 500 ms to 1500 ms, during which postsynaptic spikes were counted and binned. All simulations were done for 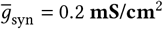 and *E*_syn_ = −45 mV.

| $N_{\text{bin}}$ | KNN | SGD | RF | SVC | GB |
| --- | --- | --- | --- | --- | --- |
| 1 | 0.612 | 0.728 | 0.746 | 0.739 | 0.746 |
| 2 | 0.658 | 0.727 | 0.759 | 0.754 | 0.756 |
| 5 | 0.770 | 0.746 | 0.782 | 0.795 | 0.788 |
| 10 | 0.725 | 0.727 | 0.793 | 0.799 | 0.806 |
| 20 | 0.670 | 0.716 | 0.778 | 0.797 | 0.806 |
| 40 | 0.596 | 0.686 | 0.772 | 0.778 | 0.805 |

To examine how classification performance improves with training set size, we repeated the ML calculations using different numbers of data points. Table 3 summarizes the variation in accuracy for classifying microcircuit motifs as the number of pulses (*N*_pulses_) increases. Each dataset contains 2*N*_pulses_ data points, of which 80% are used for training and 20% for testing under 5-fold cross-validation and mean classification accuracies are reported. For large training sets, GB achieves the highest accuracies of 0.802 and 0.806 for *N*_pulses_ = 500 and 1000, respectively. For GB with *N*_bin_ = 20, we extended the training to a larger dataset of 2000/3000/4000 points and obtained an improved accuracies of 0.817/0.824/0.832. These results suggest that, as long as the overlapping fraction of features between the two classes is not uniformly distributed, additional training data can enhance the performance of ML classifiers. We note in passing that if the external current spans a broader range than [0, 1] *μ*A in steps of 0.2 *μ*A, or is drawn from more strongly fluctuating windows, the resulting spike-count features can become increasingly variable across samples, which may adversely affect classification accuracy.

**Table 3.** Classification scores for various machine learning methods for predicting the labels of two microcircuit motifs using spike counts of postsynaptic action potential. For each microcircuit motif, *N*_pulses_ simulations were performed with random current pulses. The total simulation time was 2000 ms, and the external current was applied to the presynaptic neuron from 500 ms to 1500 ms, during which spikes were counted and binned for a time interval of 50 ms (i.e. *N*_bin_ = 20). All simulations were done for 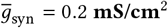 and *E*_syn_ = −45 mV.

| $N_{\text{pulses}}$ | KNN | SGD | RF | SVC | GB |
| --- | --- | --- | --- | --- | --- |
| 100 | 0.515 | 0.640 | 0.575 | 0.690 | 0.655 |
| 200 | 0.590 | 0.660 | 0.680 | 0.728 | 0.762 |
| 300 | 0.612 | 0.668 | 0.720 | 0.757 | 0.762 |
| 400 | 0.614 | 0.680 | 0.755 | 0.770 | 0.798 |
| 500 | 0.612 | 0.684 | 0.750 | 0.798 | 0.802 |
| 1000 | 0.670 | 0.716 | 0.778 | 0.797 | 0.806 |

Figure 3 shows the variation in GB classification accuracy across the parameter space spanned by the synaptic reversal potential *E*_syn_ (ranging from −20 to −70 mV in steps of 5 mV) and the maximal synaptic conductance 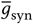 (ranging from 0 to 1 mS/cm^2^ in steps of 0.1 mS/cm^2^). We sampled 11 discrete values for each parameter, resulting in a total of 121 parameter combinations. For each combination, 100 random current pulses were applied over a duration of 1 s, with pulse widths of 25 ms and amplitudes drawn from the range 0.0–1.0 *μ*A in steps of 0.2 *μ*A (top-left panel of Figure 3). Similar calculations were repeated for narrower input ranges: 0.2–1.0 *μ*A, 0.4–1.0 *μ*A, and 0.6–1.0 *μ*A.

For sufficiently strong synaptic coupling 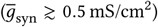, the classification behavior becomes sharply separated by the synaptic reversal potential. In this regime, the classification accuracy is predominantly either 1.0 or ∼ 0.5, with accuracies approaching 1.0 for *E*_syn_ ≳ −50 mV and collapsing to chance level for *E*_syn_ ≲ −50 mV. An accuracy of 1.0 indicates that the feedforward and feedback motifs are perfectly distinguishable based on postsynaptic spikecount features, whereas a score of 0.5 reflects indistinguishable dynamics across motifs.

At lower synaptic conductances 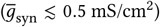, the parameter space exhibits a separating region between these two regimes, within which intermediate classification accuracies are observed. In this intermediate regime, partial overlap between spike-count feature distributions leads to scores between 0.5 and 1.0. This overlap is accompanied by a reduction in the mean postsynaptic spike counts in both motifs, with the difference between feedforward and feedback responses becoming less pronounced.

As the range of external current amplitudes is progressively narrowed, this intermediate regime becomes less prevalent. This reduction is primarily statistical in origin: drawing random current pulses from a narrower interval increases the likelihood of generating similar input realizations across samples, which in turn reduces variability in the resulting spike-count features and biases classification outcomes toward the limiting scores of 0.5 or 1.0.

## 4 Conclusions

The difference between the synaptic reversal potential and the membrane potential determines the postsynaptic response to a synaptic current. Under certain conditions, such as brain development, epilepsy, or schizophrenia, the classical distinction between excitatory and inhibitory synapses can be blurred. In this work, we constructed a map in the parameter space spanned by *E*_syn_ and *g*_syn_ to identify regions associated with distinct synaptic regimes. To this end, we simulated Hodgkin–Huxley-type equations for prototypical two-neuron microcircuit motifs under current-pulse stimulation. The binned spike counts of postsynaptic neurons were then used as features to classify the motifs with various supervised ML models. While the total spike count over the stimulus duration (1 s) provided a useful measure of average firing rate and interpretability, some methods (e.g., RF, SVC, GB) showed improved performance when using binned spike trains that capture temporal trends in firing rates. However, excessively fine binning introduced noise, which degraded classifier performance in a method-dependent manner.

This study highlights parameter regions in which excitatory (classification score ∼ 1), inhibitory (score ∼ 0.5), and ambiguous synaptic behaviors can be identified. The purpose was not to classify networks at every level of excitation or inhibition, but rather to chart distinguishable versus indistinguishable regimes in a datadriven fashion. A key challenge for machine learning methods is to improve classification performance in these ambiguous intermediate regions, where synaptic polarity is diminished and the separation between regimes is blurred.

Several limitations remain. First, the model is restricted to two fast-spiking cortical neurons, which cannot capture the collective dynamics of larger, heterogeneous neuronal populations. In particular, the present framework does not address excitation-inhibition (E–I) balance in the sense of dense recurrent networks, where irregular and noisy firing emerges from a dynamic cancellation of excitation and inhibition at the population level, as described in both theoretical models and in vivo recordings [36, 37]. In such networks, spike-train correlations and population-level statistics are used to assess synchrony and asynchrony, and the observable spike activity primarily reflects the net E-I balance, thereby obscuring the individual contributions of excitatory and inhibitory synaptic strengths. In contrast, the present work focuses on minimal two-neuron microcircuit motifs, in which synaptic strength and polarity are explicitly controlled through continuous parameters such as *E*_syn_ and 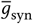. Supervised ML methods are employed to classify feedforward and feedback motifs using binned postsynaptic spike counts as features. Within this reduced setting, we identify an intermediate regime of synaptic reversal potentials in which postsynaptic spike activity becomes indistinguishable across motifs, highlighting a distinct mechanism of ambiguity that arises at the level of synaptic parametrization rather than population-level balance.

Second, only feedforward and feedback motifs were considered, whereas real neural networks also include lateral excitation, inhibition, and disinhibition motifs. Extending the framework to such motifs and to larger, heterogeneous networks would add biological realism and provide a stronger basis for mapping synaptic alterations in complex circuits. Overall, this work represents a step toward quantitatively mapping synaptic regimes in simplified microcircuits, with potential applications to understanding brain networks where excitation and inhibition are not strictly dichotomous.

## Acknowledgments

We acknowledge the support of the Department of Atomic Energy, Government of India, under Project Identification No. RTI 4007.

## Author’s Note

In previous versions of this article (v1: bioRxiv, 20 September 2025; v2: bioRxiv, 28 November 2025), a coding bug in the numerical implementation of the synaptic current led to spurious postsynaptic activity, despite the analytical expression for the synaptic current (Eq. 9) correctly stated. After fixing this implementation error, the results are fully consistent with the expected inhibitory behavior.

In particular, (i) as the range of external current amplitudes is progressively restricted from 0.0–1.0 *μ*A to 0.2–1.0 *μ*A, to 0.4–1.0 *μ*A, and finally to 0.6–1.0 *μ*A, the spike-count samples obtained for each microcircuit motif become increasingly similar. This behavior arises because the simulations are driven by random current pulses drawn from progressively narrower intervals over a fixed time window, which increases the probability of generating comparable input realizations across motifs. Consequently, the ML classification scores increasingly concentrate near the limiting values of 0.5 or 1.0, while intermediate scores become less prevalent; (ii) At *E*_syn_ = −50 mV, corresponding to a depolarizing but functionally inhibitory (shunting) regime, both feedforward and feedback motifs exhibit diminished, and in most cases absent, postsynaptic spiking activity, depending on the applied external current.

All figures and tables in the manuscript have been updated to reflect the corrected simulations. The revised ML classification scores show a slight quantitative improvement but do not lead to any qualitative change in the interpretation or conclusions.

These corrections do not alter the conclusions of the paper, namely, that the combined parameter space of *E*_syn_ and 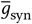 contains indistinguishable domains in which classification of microcircuit motifs becomes challenging but they improve the physical consistency and statistical clarity of the results.

